# A deep learning approach to detect and visualise sexual dimorphism in monomorphic species

**DOI:** 10.1101/2024.10.16.618646

**Authors:** Nicolas J. Silva, André C. Ferreira, Liliana R. Silva, Samuel Perret, Sonia Tieo, Julien P. Renoult, Rita Covas, Claire Doutrelant

## Abstract

Sex recognition is facilitated by dimorphism in some traits. However, humans often fail to find the traits that allow to distinguish between sexes in other species. Deep learning has the potential to surpass humans in identifying cryptic differences between sexes, but, so far, has rarely been used to assess sexual dimorphism. In this study, we evaluated (i) the ability of a fine-tuned classification neural network, EfficientNet, to find differences between sexes in a species that appears monomorphic to humans, the sociable weaver (Philetairus socius). We then assessed (ii) the benefits of Grad-CAM visualisation techniques to understand which parts of the individuals are used by the network to differentiate the sexes. We trained 10-folds cross-validation models on more than 4,500 pictures of the head from more than 1,300 individuals. Our results show that the network can predict sex of sociable weavers with an accuracy of 76%, which is considerably higher than humans’ performance, and that the model was similarly good at predicting females and males. When interpreting the probability of being classified to one sex, our results further reveal an effect of the interaction of sex with age on the confidence score of the models which shows that younger males are less masculine than older ones, and older females more masculine than younger ones. Finally, using Grad-CAM we found that the model mostly uses the beak region to predict the sex of individuals. Overall, this work shows that artificial intelligence has the potential to be a non-invasive sexing tool, surpassing human capabilities and aiding in pinpointing potential cryptic dimorphic body parts that have yet to be identified. In birds, half of the world’s species appear sexually monomorphic to humans, and re-evaluation of species dimorphism with this type of methods could deepen our understanding of the effect of selection on animal traits.

## 1. Introduction

Sexual dimorphism can be present in few or several species’ traits. It is the result of selection on those traits (in particularly sexual selection and predation), drift and phylogenetic trajectories (Badyaev & Hill, 2003). Exploring sexual dimorphism in different species allows to determine how the evolutionary pressures may vary within and between the sexes. Sexual dimorphism is usually based on differences on size, shape or colouration (Hedrick & Temeles, 1989; Corl et al., 2010; Warren et al., 2013; Kuntner & Coddington, 2020; Mori et al., 2022), and is common in taxa such as birds (Andersson, 1982), fish (Kodric-Brown & Brown, 1984), mammals (Caro, 2009) and invertebrates (Allen et al., 2011). Humans, usually assess these traits through direct observations of the visual aspect of the individuals (Badyaev & Hill, 2000; 2003). However, our ability to detect dimorphism is limited by the capacity of our visual system that may differ from the ones of animals, and often only the most conspicuous dimorphism can be reliably identified by human observers. This limits our ability to detect different types of dimorphism, and thus to study their adaptive value (Villafuerte & Negro, 1998; Hung et al., 2017; Ó Marcaigh et al., 2021).

Different approaches have been developed to overcome the issue of detecting sexual dimorphism. For instance, Eaton (2005) used spectrophotometry measurements of reflectance and, considering birds’ colour discriminatory abilities between 300nm and 700nm, showed that on 139 presumed sexually monochromatic bird species from the Passeriforme order, more than 90% were dichromatic from an avian visual perspective. Similar measures were used in a jumping spider (*Phintella vittata*) revealing a UV-B sexual dimorphism in this species (Li et al., 2008). More recently, computer vision has been shown to be effective in measuring colouration. For instance, Support Vector Machine, a machine learning tool, was used to classify sex of zebrafish (*Danio rerio*) based on caudal fin colouration (Hosseini et al., 2019). Support Vector Machine requires human interference to pre-define which features to learn: i.e., to identify and measure the specific visual features that allow to distinguish and classify sexes. Knowing the sexually dimorphic trait before the analyses has the main drawbacks of potentially missing important parts and features of the animals that might be useful for sex classification by the animals, but are cryptic to humans.

Deep learning methods, and in particularly Convolutional Neural Networks (CNNs), do not require predefinition of the relevant features and can overcome this limitation by automatically determining the features that are optimal to perform a given classification task (Jordan & Mitchell, 2015; Angermueller et al., 2016; Christin et al., 2019; Chollet, 2021). Solving category classification problems using CNNs is now increasingly used in the fields of ecology and evolution (e.g., Christin et al., 2019; Ferreira et al., 2020; Hansen et al., 2020; Norman et al., 2023). Specifically, CNNs have been previously used in species with known sexually dimorphic traits in order to increase processing speed and efficiency of sex recognition, but to date, mostly in humans (Dwivedi & Singh, 2019). For instance, deep learning was used on dentomaxillofacial features extracted from radiographs to sex children at an early age, when sexual dimorphism is not well marked yet (Franco et al., 2022). Similarly, on human skulls, a deep learning network analysing computed tomography scans of human brains was able to classify women and men with an accuracy of 95% (Bewes et al., 2019). In animals, Chen et al. (2021) showed that deep learning can be used to classify females and males in 30 *Pseudopoda* spider species based on their genitals (Chen et al., 2021); and Hosseini et al. (2019) also used CNNs successfully to classify sex based on colour on zebrafish pictures. However, CNNs have been rarely used to find sexual dimorphism in traits that appear monomorphic to the human eye, but in the gray wolf (*Canis lupus*) deep learning revealed a previously unnoticed sexual dimorphism in cranial shape (MacLeod & Horwitz, 2020). In *Pseudopoda* spider species, Chen et al. (2021) also used deep learning identify sex-specific patterns and colours of the back of individuals (that are sexually monomorphic for humans) and found sexual dimorphism in 94% of their dataset. Finaly, a recent study on giant pandas (*Ailuropoda melanoleuca*) faces succeeded to perform sex classification with an accuracy of 77% using CNNs (Wang et al., 2019).

A major drawback of using CNNs to classify the sex of animals, however, is that the features automatically built and used by the algorithm for performing classification are unknown (without additional analyses), or are difficult to represent in a way that is meaningful for humans (i.e., CNNs work as ‘black boxes’, see Alain & Bengio, 2016; Castelvecchi, 2016; Shwartz-Ziv & Tishby, 2017; Lei et al., 2018). In order to overcome these limitations, tools have been developed to understand what do CNNs learn. In computer vision, one of the most widely used tools is the Gradient-weighted Class Activation Mapping (Grad-CAM) (Selvaraju et al., 2017).

Grad-CAM creates an activation thermal map for a specific class that is superimposed with the original image. Every channel of the picture (i.e., the different dimensions of an image pixels) is weighted with the class gradient of the channel. These Grad-CAMs colour the pictures to identify which parts of the images were the most relevant for classification, and thus the possibly most important features to discriminate sex. For instance, in giant pandas, Grad-CAMs qualitatively showed sex difference in the eyes and in the nose of the individuals (Wang et al., 2019).

Here, we developed a deep learning tool based on EfficientNet (Tan & Le, 2019) and used a monomorphic bird species, the sociable weaver (*Philetairus socius*) (MacLean, 1973) to (i) explore whether species that appear sexually monomorphic to humans can be detected as sexually dimorphic by CNNs based on pictures. In addition, (ii) we explored the use of Grad-CAM visualisation techniques to pinpoint which parts of the individuals are used by the network to differentiate the sexes. In birds, it is common to observe delayed plumage maturation (Hawkins et al., 2012), making young males generally more look alike females. We therefore (iii) tested whether the accuracy of sex identification varied with age, expecting younger males to be accurately classified less often than older males.

## 2. Material and Methods

### 2.1. Species and data collection

We worked with a population of sociable weavers inhabiting Benfontein Nature Reserve, in South Africa (−28°818S, 24°816E, 1190m asl) and monitored since 1993 (Covas et al., 2002). Since 2008, birds in the study colonies are caught every year, usually at the end of winter (August-September). During, the capture events, a blood sample is taken and used for genetic sexing (Covas et al., 2006). Additionally, top view pictures of the head of the birds are taken following an established protocol (Rat et al., 2015).

For most of the birds, breeding monitoring allows us to know their exact age. However, individuals can move between colonies (especially females; Covas et al., 2006; van Dijk et al., 2015). For immigrant birds first caught as adults in the studied population, we consider them to have the average population dispersion age at capture. We computed this age by adding to the first date of capture the average minimum age of first dispersion measured in the population (690 days for males and 727 days for females to the date of first capture (Silva et al., in prep)).

### 2.2. Ethical note

Birds were captured and handled using protocols approved by the Northern Cape Nature Conservation (permit FAUNA 1638/2015, 0825/2016, 0212/2017, 0684/2019 and 0059/2021) and the Ethics Committee of the University of Cape Town (permit 2014/V1/RC and 2018/V20/RC). The birds were captured using mist nests and were extracted directly after being caught. All efforts have been made to minimise handling time of the birds (i.e., task division of the processing steps of extracting the birds from the mist nests, ringing the birds, taking blood samples, taking pictures; release of the birds was closely monitored to detect any uncommon flight behaviour). These procedures have been performed during the non-breeding season to avoid disturbing birds with chicks at the nests. In total, 1,323 caught individuals were included in this study (see more details in the following subsection).

### 2.3. Datasets

We obtained 4,595 images (2,246 for females and 2,349 for males) depicting 1,323 individuals (661 females and 662 males) photographed during six annual capture events (between August 2015 and Septembre 2021 - no captures in 2020). We selected only individuals with a fully developed adult plumage. At each capture event, one to four pictures per individual were collected ((mean±SD) females: 2.63±0.81; males: 2.46±0.88). We trained and used a Mask-RCNN model to extract the birds’ heads from full pictures (see supplementary 1 for description of the model). All pictures were orientated with the throat of the individuals facing the bottom of the pictures, in a portrait style (Fig. 1.A). On the 1,323 individuals in our dataset, 993 were assigned with their real age ((mean±SD) 773±560 days old) and 330 with an estimated age ((mean±SD) 1,372±924 days old).

**Figure 1:**
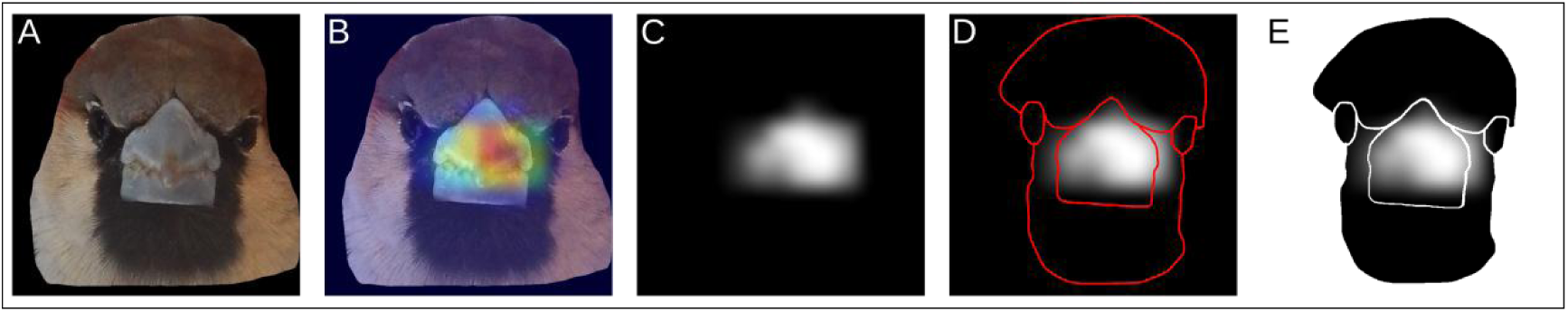
Workflow of picture processing along the training and visualisation process. (A) The picture is segmented from a bigger picture with a first neural network to keep only the head of the individuals with their bib facing down. (B) Blue to red scale Grad-CAM superimposed with transparency to the original picture. It allows us to visualise which regions of an image the model is using to perform its classification. (C) Grey scale Grad-CAM. (D) Grey scale Grad-CAM with the shape of the regions of interest of the head of the birds: the cap, the beak, the bib and the eyes. (E) Each region is exported on its own and analysed separately.

**Figure 2:**
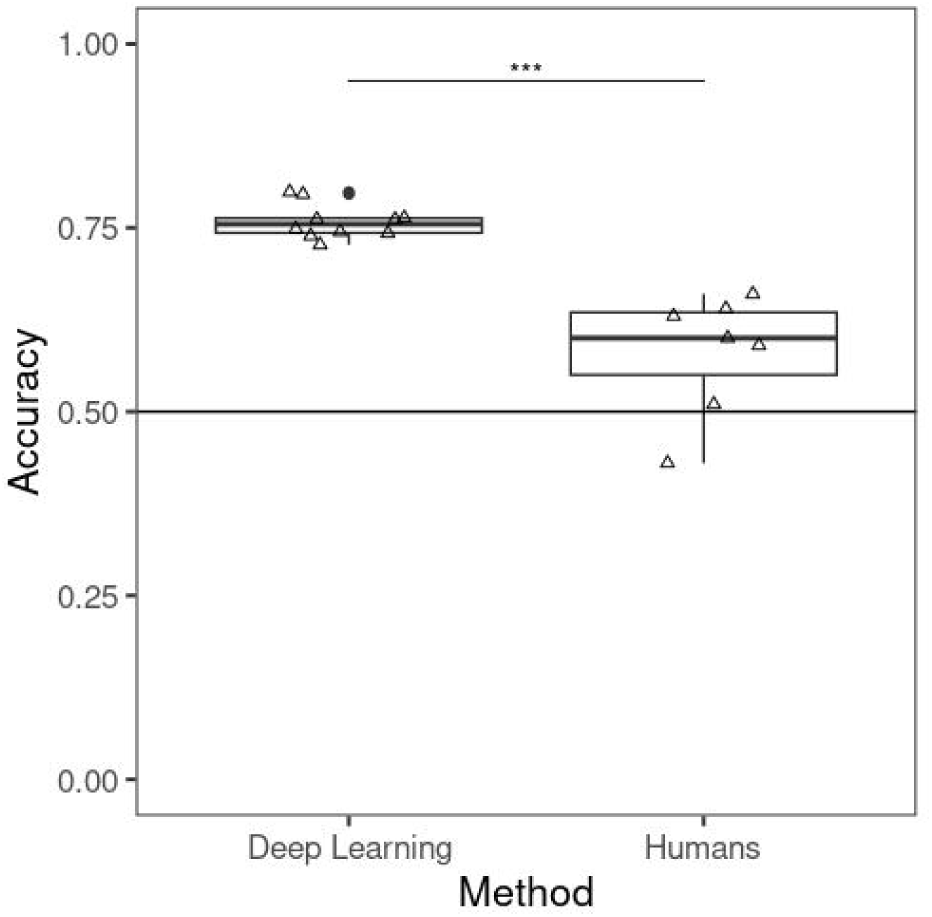
Accuracy of the individual classification in males and females in function of the classification method used. The deep learning accuracies were obtained from the predictions on the test dataset at each of the 10-folds during the training. The human accuracies were obtained from blind visual classification by humans. The classification methods were found to be significantly different from each other and they were both found to be significantly different from random classification. Triangles represent raw data.

### 2.4. Model training

We used transfer learning from an EfficientNet B4 network (Tan & Le, 2019) pre-trained on ImageNet database (Deng et al., 2009) composed of more than 14 million pictures to classify objects from images. This method allowed us to use the weights of an already trained model and fine-tune them for the task we desire. Since our objective is also image classification (i.e., classify images as males or females), this pre-trained model on ImageNet network is an appropriate choice to perform transfer learning. During the fine-tuning of the network, we implemented average pooling throughout the model’s backbone to prevent over-fitting during the training. We replaced the fully connected part at the end of the EfficientNet B4 network with a dropout layer (rate: 0.4), a dense layer with 1,024 neurons and a ReLU activation, a dense layer with 64 neurons and a ReLU activation, and a final dense layer with one neuron and a Sigmoid activation. The dropout layer was added to limit over-fitting of the model by randomly setting to zero some neurons of the network (Srivastava et al., 2014). We compiled the model with an Adam optimizer (Kingma & Ba, 2014), a binary cross-entropy loss and a binary accuracy. All layers were set as trainable except for the BatchNormalisation layers. Due to the sigmoid last layer, the model outputs are numbers between zero and one, with [0; 0.5 [being scores of sex prediction to females and [0.5; 1] being scores of sex prediction to males. To better compare males and females, we reversed the females’ score (one minus the predicted score). Therefore, for all individuals, a final score between [0; 0.5 [signal an erroneous sex prediction and a score between [0.5; 1] a correct sex prediction.

The hyper-parameters of the training process were inferred by training a preliminary model using 90% of the individuals (595 males, 596 females, 4183 pictures) for training dataset and 10% for validation (66 males, 66 females, 412 pictures). This preliminary model helped us to define the following parameters for training. The pictures were re-sized to 400×400 pixels and passed through the network with a batch size of 16 pictures. The training process was performed on 13 epochs with a learning rate of 10^-3^ from epoch one to three to quickly learn general patterns for sex classification, then learning rate was set to 10^-4^ to help the model to learn more precise features.

We then performed 10-fold cross-validation (again 90% of the individuals as training dataset and 10% of the individuals as test dataset). At each fold, individuals from the test dataset were swapped with an equivalent number of individuals from the training dataset. This allowed each individual to be in a test set once during the training, and thus to get one prediction of sex per image. We performed data augmentation and applied to the pictures random horizontal flip with a probability of 0.5 and random contrast between [-0.2; 0.2] with a probability of 0.5. Additionally, we evaluated the impact on the training process of reducing by a quarter, half and three quarters the number of individuals on in the training dataset (Table S1).

### 2.5. Model visualisation

We used Grad-CAM (Selvaraju et al., 2017) to identify the specific areas of the images that influenced the model’s decision in classifying individuals as male or female. We computed on the Grad-CAM of the last 2D convolution layer (called “top_activation”) to calculate the probability of being a male (because we used a sigmoid activation for the last layer). The different regions of the pictures were coloured from blue to red, respectively from lower to higher importance, depending on the role they had in the network’s decision (Fig. 1.B). Analysis for this part was adapted from Chollet (2021).

### 2.6. Statistical analysis

All statistical analyses were conducted on R version 4.3.1.

#### 2.6.1. Sex recognition

The average accuracy of the models on the test datasets was saved (N=10 models) at the end of each fold during the training process. These accuracies were compared to those reached by human observers. To evaluate human performance, seven of the authors (AUTHORS’ NAMES HIDDEN FOR THE REVIEWING PROCESS) were first given the same set of labelled pictures of 25 females and 25 males as an attempt to train themselves to distinguish males and females. Then, they manually and blindly classified each a 100 randomly selected images of unique individuals (50 males and 50 females). Then, we performed a first Wilcoxon test to compare the accuracies of the machine and humans.

We simulated a random classification of sex of the 100 birds classified by humans using a binomial distribution with a probability of 0.5. With Wilcoxon signed rank exact tests, we tested if the accuracy reached by the CNNs and the one reached by humans were different from a random classification.

#### 2.6.2. Visualisation of the features used for classification

We quantitatively measured which parts of the pictures the models are using to classify males and females. For each of the 10 training folds, we randomly sampled, from the test dataset, 10 pictures of males and 10 pictures of females that were rightly classified. We manually segmented these 200 pictures (Fig. 1.D) to identify four parts of interest on the head of the birds: the cap, the beak, the bib and the eyes (Fig. 1.E).

We computed Grad-CAM images with a grey scale colouration (pure black indicates no activation and pure white indicates very high activation) (Fig. 1.C). On these images, using GIMP version 2.10.36, for each region we measured, a) the proportion of activated pixels to estimate the surface proportion of each of the four regions used by the model for sex classification; and b) the average activation of the activated pixels from] 0; 1] (one indicates pure white). The first approach identifies the regions of the head that is most used by the model by considering its surface, while the second approach tells us about the importance of each activated region among the activated pixels in the pictures. Both measurements are useful to identify dimorphism because the first tells us how much of a region is used for classification and the second helps to pinpoint the intensity of a region. For instance, by relying on both measurements we could distinguish regions with a large proportion of pixels activated at low intensity from regions with only a small portion of its surface activated but with very high intensity. Because of the sigmoid last layer, we predict the probability for an individual to be a male. Therefore, high difference of activation between males and females indicates higher dimorphism on the trait, while if the activation for a trait is similar between males and females, then the traits isn’t very dimoprhic.

We performed multiple Student tests to determine if the proportion of activated pixels per regions and the average activation were different between sexes, and multiple paired Student tests to understand if these two measures were different between regions within the same individual. P-values were adjusted with the Benjamini-Hochberg procedure (Benjamini & Hochberg, 1995).

#### 2.6.3. Sex and age difference on the classification performance

We tested if males and females differed in their likelihood of being correctly identified and if age also influenced the classification performance by running two mixed models.

i. We performed a generalised linear mixed model (GLMM) (Binomial error distribution with a link logit). As response variable, we binarised the average predicted score per individual, with 1 indicating rightly classified individuals and 0 wrongly classified ones. As fixed factors, we used sex, and we added as random factor the identity of the individuals, the cross-validation fold and the year of data collection. Several pictures per individuals were used and it is important to specify in the model they are not independent points. The pictures in a test dataset during the cross-validation folds were evaluated with the same model and therefore their predicted score is not independent. Last, different people between years were involved in the picture collection of the birds, so we controlled for intra- and inter-photographer variation.
ii. We fitted a GLMM (Beta error distribution with a link logit). The response variable, was the score predicted by the model (i.e., the confidence score given by the deep learning model). This score indicates how confident the model is about its classification decision. A score closer to 1 indicates that the sex classification is relatively robust. We used sex, age, and their interaction as fixed factors, to account for the expectation of males becoming more masculine with age, an effect not anticipated for females. As random factors, we included the identity of the individuals, the cross-validation fold and the year of data collection. 27 pictures (18 males and 2 females individuals, (mean±SD) 1,009±825 days old) were predicted with a confident score of exactly 1, but because Beta distribution only accepts continuous values in range] 0; 1[, we substracted a value of 10^-10^ for the pictures to be included in the model. Age was scaled and mean-centered. Since age is either the true age or the estimated age for immigrants, the model was computed first with the individuals with the exact known age only (992 individuals, 3,445 pictures), and second with all the individuals (1,323 individuals, 4,595 pictures). The two models were similar, except for the effect of age in females, that tends to be negative in the model including all individuals and is negative in the model including only individuals with exact age (Table S2). We decided to be conservative and keep only the individuals with known exact age.

Mixed models were computed with lme4 package version 1.1-34 (Bates et al., 2014) and glmmTMB package version 1.1.8 (Brooks et al., 2017) for the GLMM Beta. Plots of models were created with ggplot2 package version 3.4.2 (Wickham, 2014), ggeffects version 1.3.1 (Lüdecke, 2018) and sjPlot package version 2.8.15 (Lüdecke, 2023).

## 3. Results

### 3.1. Sex recognition

On average, we found that the models were able to classify males and females with an average accuracy of (mean±SD) 75.8%±2.4% at the picture level within the 10 folds (Fig. 1), and very strong evidence (Wilcoxon test, V=0, P<0.001) that these classifications were different from a random guess ((i.e., accuracy=0.5). Human classifications were also found to be different from a random guess (Wilcoxon test, V=0, P<0.001) but they were less accurate (mean±SD) 58.0%±8.2% (N=7 humans; Fig. 1) than the deep learning models’ performance (Wilcoxon test, W=70, P<0.001; Fig. 1). These results show that the deep learning method is able to classify the sex of sociable weavers with higher accuracy than humans.

### 3.2. Visualisation of the features used for classification

Males presented both higher proportion of activated pixels and a higher intensity in the activated pixels than females for all of the four regions delimited in the head of the birds (beak, bib, cap and eyes: Fig. 3A,B; Table S3). Among these four regions, both the highest proportion of activated pixels and the highest intensity in the activated pixels were observed in the beak for males, and in the bib for females (Table S4). This indicates that the four regions are used by the model to differentiate males from females, but that the model focuses more intensely on the beak of the individuals to perform its classification. The fact that the bib is also activated in females may indicate that the bib is a source of misclassification and that not many sex difference can be found on that trait.

**Figure 3:**
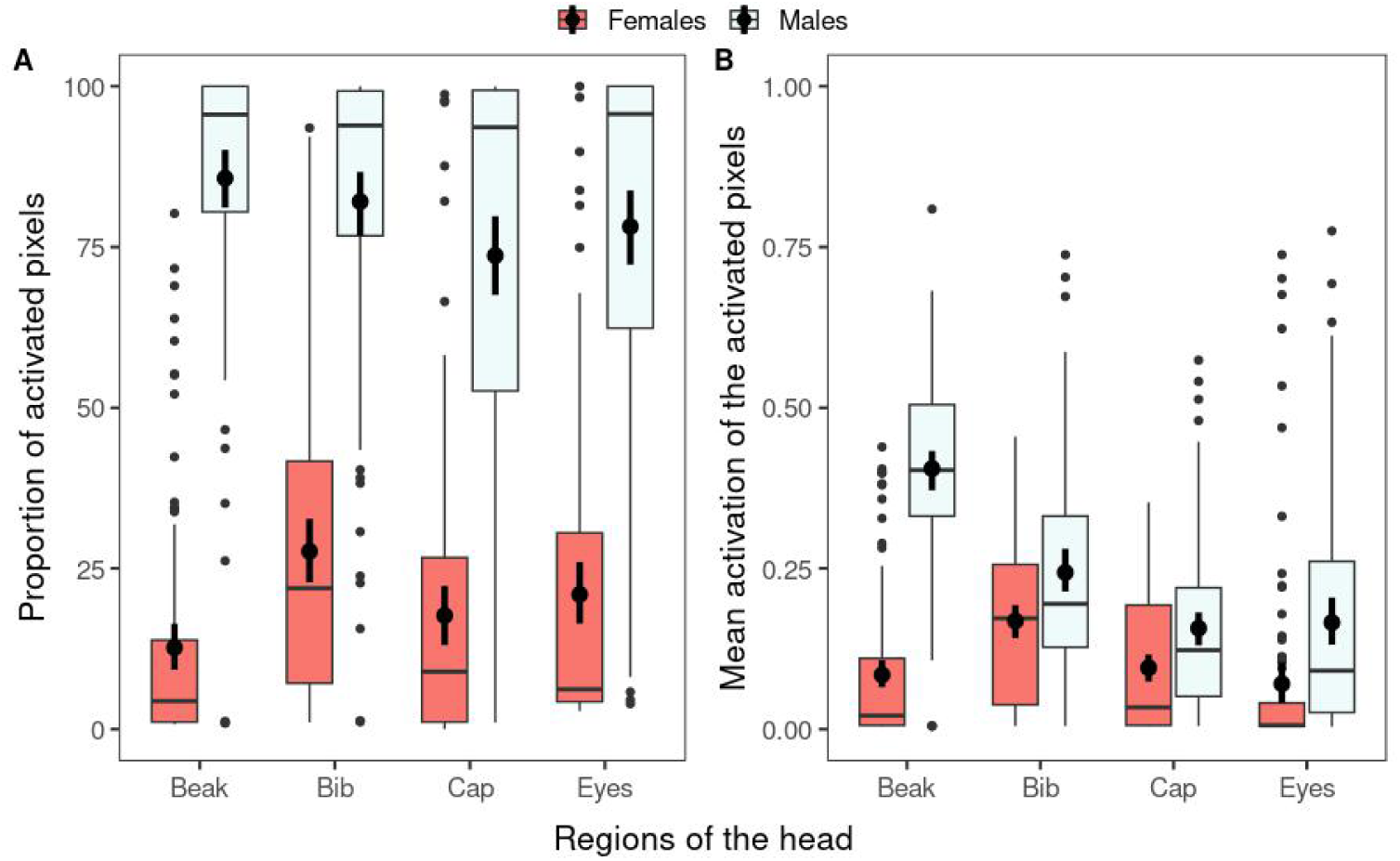
Difference in (A) proportion of activated pixels and (B) mean activation of the activated pixels, in function of sex and region of the head. Females are in red and males in light blue. The beak of the individuals seems to be the main feature used by the models to perform its sex classification (more proportion of activated pixels and higher activation intensity in males than in females). Boxplots represent median and quantiles at 25% and 75%, and error bars represent mean and 95% confidence intervals.

### 3.3. Sex and age difference on the classification performance

We found no evidence that sex has an effect on the probability to rightly classify a picture of an individual ((estimate±SE (95%CI)) −0.19±0.17 (−0.53; 0.14), z=-1.12, P=0.260; Table 1), indicating that no sex is better classified than the other one.

**Table 1:** Estimate of the GLMM parameters to test the effect of sex on the probability for the deep learning network to correctly classify sex from pictures of the individuals’ head. The model was performed on 4,593 pictures, with individuals with both the exact age and the estimated minimum age.

| Fixed effect | Estimate | SE | 95% confidence interval | z-value | p-value |
| --- | --- | --- | --- | --- | --- |
| Intercept | 2.28 | 0.15 | 1.99 ; 2.57 | 15.27 | <0.001 |
| Sex - Male | -0.19 | 0.17 | -0.53 ; 0.14 | -1.12 | 0.261 |
| Random effect | Variance |  |  |  |  |
| Individual (N=1,323) | 5.591 |  |  |  |  |
| Season (N=6) | 0.000 |  |  |  |  |
| Cross-validation fold (N=10) | 0.000 |  |  |  |  |
| Residuals | 3.290 |  |  |  |  |

We found a strong evidence for an association between predicted score and age that differed between the sexes ((estimate±SE (95%CI)) 0.43±0.07 (0.30; 0.56), z=6.47, P<0.001; Fig. S1; Table S2.B), with a positive link for males and a negative for females (Fig. 4; Table S2.B). This indicates that, for males, younger individuals are less well classified than older ones, while for females it is the reverse.

**Figure 4:**
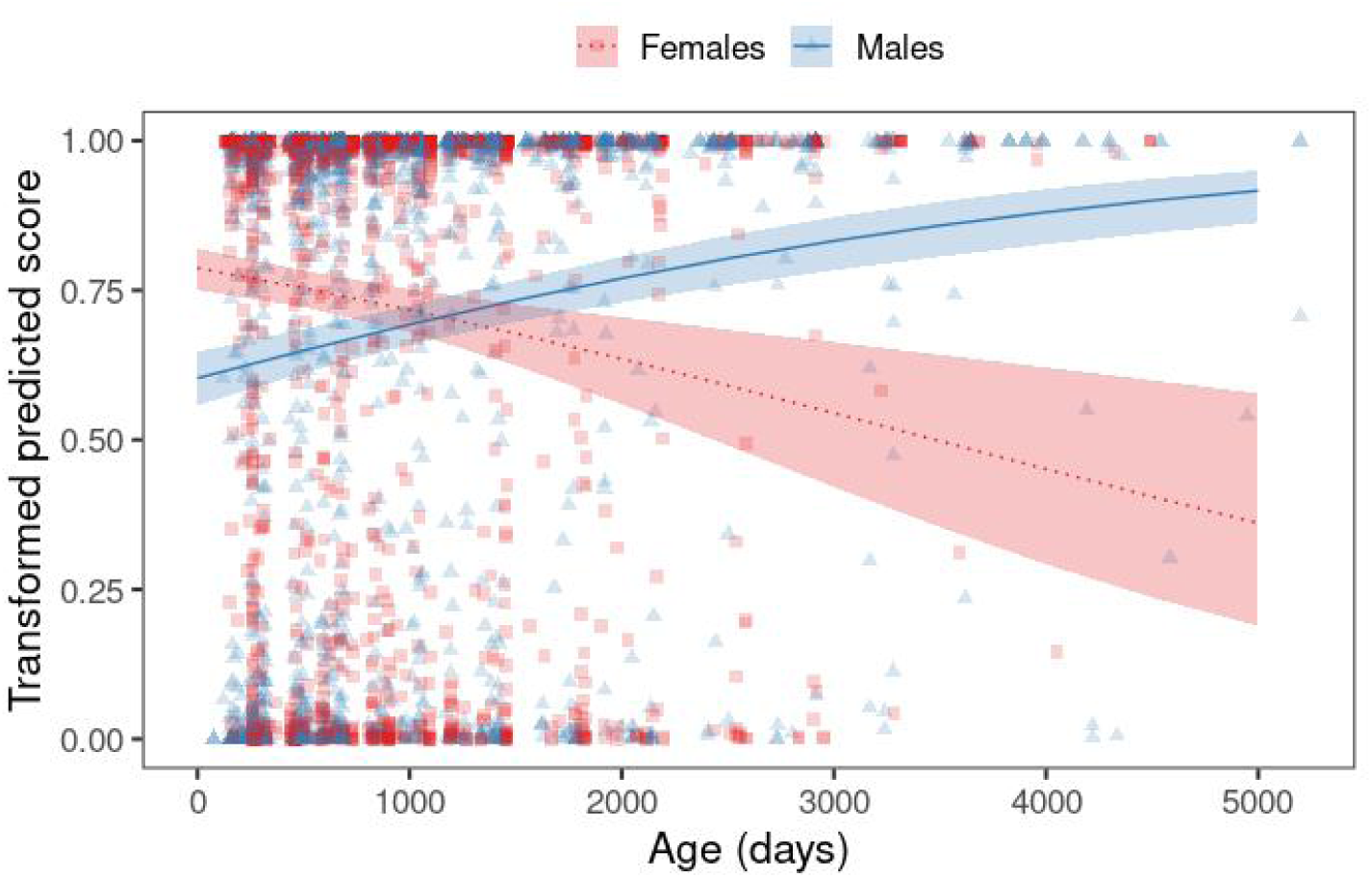
Score predicted by the deep learning model in function of age and sex. Females are represented in red with dashed line and squares, and males are represented in blue with solid line and triangles. Squares and triangles are the raw data used to fit the mixed model. Lines correspond to estimated means obtain with the mixed model and shaded area to the 95% confidence intervals. Older males are better predicted than younger males while the opposite is found for females. Female predicted scores were transform such that 1 corresponds to the best predictions for both sexes.

## 4. Discussion

Our results show that a deep learning-based method surpasses human performance when classifying the sex in a bird species with limited sexual dimorphism, and that by using Grad-CAM we were able to pinpoint differences used by the algorithm to differentiate between males and females. Finally, by investigating the source of misclassifications of the models, we found that age influenced differently females and males when predicting sex using CNNs.

Species visually monomorphic for human eyes are common, and in these cases long and costly behavioural observations or genetic methods are necessary to determine the sex of individuals. For instance, in birds, half of the world species does not show clear differences between males and females for human eyes (Griffiths et al., 1998). If the CNNs identify features that are detectable for animals, this will have implications that will allow to better understand how sexual and natural selection may have shaped the evolution of these species’ morphological traits. Deep learning methods require large volumes of data to be trained and thus time and funding to collect these data. In addition, running deep learning models use large amounts of energy. However, once a model is trained it can be indefinitely re-used, improved and can often outperformed humans (e.g., Ditria et al., 2020; Ouyang et al., 2020; Jocher et al., 2022; this study).

Here, we reached only 76% accuracy, similarly to the one obtained on giant panda faces (Wang et al., 2016). However, we only used a small portion of the bird’s body and future work could potentially increase model’s performance by considering, for instance, photos from multiple body parts of the birds at the same time. In addition, increasing the training sample size and including more pictures of more individuals at different ages is also expected to improve model accuracy. Our results (Table S1) show that the efficiency of our trained network decreased when decreasing the training dataset sample size, indicating that we may not have reach the full potential of using CNNs for sex classification in our species. Here, we selected very standardised pictures to train and test our models, and further tests could be performed with pictures taken under more natural conditions (e.g., at bird feeders; see in Ferreira et al., 2020). Hence, we suggest that in some monomorphic species and with large enough training datasets, and photos from different parts of the body, sexual recognition of individuals using deep learning models could potentially achieve sufficient performance to replace genetic sexing, which, in the long run, could save time and research funds. Importantly, these methods have the potential to limit animal suffering, pictures can be taken without capturing individuals (e.g., Ferreira et al., 2020) and avoiding the capture and handling stress.

With the Grad-CAMs, we were able to find a sexual dimorphism on the beak of the individuals. No sexual differences based on the head of the individuals were ever determined based on morphological measurements or the field obervations, and the sexes appeared undistinguishable (Maclean, 1973; Covas & Doutrelant, unpublished data). Sexual dimorphism based on the beak of individuals have been described in other weaver species (e.g., red-billed quelea (*Quelea quelea*) (Walsh et al., 2012); red-billed buffalo weaver (*Bubalornis niger*) (Maclean, 1993); african village weaverbird (*Ploceus cucullatus*) (Collias & Collias, 1970); white-browed sparrow-weaver (*Plocepasser mahali*) (Leitner et al., 2009)). However, the dimorphism described for these species, based on bill length and/or colouration, is visible using measurements, which in not the case for the sociable weavers. Further investigation is now necessary to understand which features are used by the model used here to classify sex (e.g., shapes, colours, contrasts). The mechanisms used by the deep learning models to identify the critical features is usually treated as a ‘black box’ (e.g., Alain & Bengio, 2016; Castelvecchi, 2016; Shwartz-Ziv & Tishby, 2017; Lei et al., 2018). Grad-CAMs are one of the first steps to overcome this issue and understand potential cryptic differences between categories. In dimorphic species, many studies have investigated the potential adaptive value of conspicuous traits (e.g., colouration, size), linking them to fitness benefits (e.g., reproductive success, mating success). Our results highlight the potential to investigate similar questions on dimorphic traits which are cryptic to the human eye, but can be detected by a deep learning neural network, that could potentially be under the same evolutionary pressures as conspicuous traits.

When dealing with classification problems, particularly in the field of computer science, there is a push to maximise performance metrics (e.g., to obtain nearly 100% accuracy). However, in the biological sciences, inefficiency of the neural networks could also inform us about cryptic features of animals (or organs, cells, etc.) that might open the door to new, biologically relevant hypotheses. For instance, deep learning was used in mandrills (*Mandrillus sphinx*) to assess face similarity of individuals, revealing higher similarity among kin and allowing to explore spatial associations between similar-looking individuals (Charpentier et al., 2022). Here, we took advantage of the imperfection of our models regarding sex determination to reveal that young males are less masculine than older ones and that older females look more masculine than older ones, something that as been already observed in primates (Kloth et al., 2015; Tieo et al., 2023). In birds, however, only a lower masculinity of younger males has been regularly observed with notably plumage delayed maturation (Hawkins et al., 2012), while, to our knowledge, no quantitative description of female masculinisation with age has been reported in a non-primate species (although some descriptions exist, for example older females gang-gang cockatoos (*Callocephalon fimbriatum*) can acquire the orange colouration in the head that is typical of males; Forshaw & Cooper, 2002). This is not surprisingly, particularly in apparently monomorphic species, where humans cannot directly obtain information about which regions of the individuals are attributed to a masculine or a feminine appearance, hindering attempts of investigating how they change with age. We note here that for very old female sociable weavers, the relationship between predicted score and age could arise from the male bias sample size in the training dataset (0 females and 21 males older than 3,000 days; Fig. S2). However, removing these old individuals from the dataset used for the mixed models did not change the observed relationship for younger individuals, although with a slight change in effect size (Table S5). This indicates that despite the unbalanced sex ratio of very old individuals, we can still be confident, for most individuals, of the relationship between predicted score and age in relation to sex that we found. Our approach and results indicate that in birds, similarly to primates (Charpentier et al., 2022; Tieo et al., 2023), deep learning could be used to study how sex and age (or other relevant traits) interact and affects the visual aspect of individuals. In addition, this also indicates that potential features expressing age may be present in the head of the individuals (e.g., age classification of giant pandas (Qi et al., (2022)). These results open the door to address the potential adaptative value of such interactions.

## 5. Conclusion

Our study emphasises the potential for deep learning methods to surpass human abilities in detecting sexual dimorphism in species. It also helped us to identify the beak as a trait potentially indicating sex in our study species. We hope that this approach will stimulate interest among researchers to re-assess sexual monomorphism in their study species, and how it varies in relation to individual attributes. Sexual dimorphism is one of the main traits used in sexual selection studies and better quantification of species dimorphism could influence our understanding of the selective forces acting in each focal species, potentially uncovering new avenues for research.

## Acknowledgments

We thank all the people that helped in field data collection along the years and in particular Annick Lucas and Margaux Rat. We also thank Franck Théron for the management of the long-term database. Access to Benfontein Nature Reserve was provided by De Beers Mining Corporation. This study was supported by funding from ANR to C.D. (France, grant 19-CE02-0014-01), ERC to R.C. (EU, Consolidator grant 866489) and by the DST-NRF Centre of Excellence at the Fitzpatrick Institute of African Ornithology University of Cape Town. N.J.S. was funded by the ANR 19-CE02-0014-01, L.R.S by ERC-Consolidator 866489, R.C. by FCT (CEECIND/03451/2018) and A.C.F by University of Zurich Forschungskredit postdoc grant (K-74312-01-01 University of Zurich), Swiss Federal Commission for Scholarships and by European Research Council (grant agreement no. 850859 awarded to Damien Farine). This project is part of the OSU OREME, and long-term Studies in Ecology and Evolution (SEE-Life) program of the CNRS.

## Author’s contribution

N.J.S., A.C.F., L.R.S., R.C. and C.D. conceived the study. N.J.S., A.C.F, L.R.S., S.P., R.C. and C.D. collected long-term data. N.J.S., A.C.F., L.R.S., S.P. pre-processed the data. N.J.S., with the input from A.C.F., L.R.S., S.T. and J.P.R., led the deep learning model training and the statistical analyses. N.J.S led the writing of the manuscript. R.C. and C.D. acquired the funding and led the long-term program that made this work possible. All authors contributed critically to the drafts and gave final approval for publication.

## Data availability

Data are available in the Dryad repository : https://datadryad.org/stash/share/hZ8_tpL2XtIzvEH3pDBT4HsF9EExA2sdVb22foe7oh8

