## Supplementary materials for "A deep learning approach to detect and visualise sexual dimorphism in monomorphic species"

[1] Rita Covas and Claire Doutrelant should be considered joint last authors.

### Supplementary materials

**Appendix 1: Mask-RCNN model to segment and extract the birds' heads from pictures.**

A mask-RCNN model was trained to automatically extract the head of the birds from the background (following: https://github.com/matterport/Mask_RCNN). We used a ResNet101 backbone pre-trained on Coco dataset (Lin et al., 2014). We used 427 pictures to train the model (316 for training dataset and 111 for validation dataset), for which we manually delimited the silhouette of the birds' heads. We stopped training after 20 epochs, having qualitatively verified that the model successfully extracted the heads of the birds from the validation dataset.

**Appendix 2: Effect of the size of the training dataset on model accuracy.**

We evaluated the influence of reducing the size of the training dataset on the accuracy of sex classification by the model. We randomly removed one third, one half, and three third of the individuals present in the training dataset. We performed multiple Student tests to determine if model accuracies were different. P-values were adjusted with the Benjamini-Hochberg procedure (Benjamini & Hochberg, 1995). We found that reducing sample size also reduced the global accuracy of the models (Table S1).

**Table S1:** Effect of reducing the size of the training dataset on the accuracy of sex classification. Data were obtained using 10-fold cross-validation and the same architecture, detailed in the Materials and methods section. P-values were adjusted with the Benjamini-Hochberg procedure (Benjamini & Hochberg, 1995).

| Percentage of training dataset | Statistic | Adjusted p-value |
|---|---|---|
| 100 vs 75 | 2.31 | 0.033 |
| 100 vs 50 | 6.21 | <0.001 |
| 100 vs 25 | 8.21 | <0.001 |
| 75 vs 50 | 4.02 | 0.001 |
| 75 vs 25 | 6.42 | <0.001 |
| 50 vs 25 | 3.14 | 0.007 |

#### Appendix 3: Sex and age difference on the classification performance

**Table S2:** Estimate of the GLMM parameters to test the effect of sex and age on the score predicted by the deep learning network from pictures of the individuals' head. Model A was computed with all the individuals in the dataset (N=4,593 pictures), and model B was computed with only the individuals with exact known age (N=3,463 pictures). Conditional $R^2$ of model A and model B are respectively of 0.91 and 0.93.

**Model A**

| **Fixed effect** | **Estimate** | **SE** | **95% confidence interval** | **z-value** | **p-value** |
|---|---|---|---|---|---|
| **Intercept** | 1.02 | 0.07 | 0.87 ; 1.16 | 13.92 | <0.001 |
| **Age** | -0.08 | 0.04 | -0.16 ; 0.01 | -1.79 | 0.073 |
| **Sex** - *Male* | -0.22 | 0.06 | -0.34 ; -0.11 | -3.80 | <0.001 |
| **Age** : **Sex** - *Male* | 0.31 | 0.06 | 0.21 ; 0.42 | 5.67 | <0.001 |

| **Random effect** | **Variance** |
|---|---|
| Individual (N=1,323) | 0.670 |
| Season (N=6) | 0.013 |
| Cross-validation fold (N=10) | 0.012 |
| Residuals | 0.053 |

**Model B**

| **Fixed effect** | **Estimate** | **SE** | **95% confidence interval** | **z-value** | **p-value** |
|---|---|---|---|---|---|
| **Intercept** | 1.02 | 0.08 | 0.85 ; 1.18 | 12.14 | <0.001 |
| **Age** | -0.21 | 0.06 | -0.32 ; -0.10 | -3.68 | <0.001 |
| **Sex** - *Male* | -0.29 | 0.07 | -0.42 ; -0.16 | -4.34 | <0.001 |
| **Age** : **Sex** - *Male* | 0.43 | 0.07 | 0.30 ; 0.56 | 6.47 | <0.001 |

| **Random effect** | **Variance** |
|---|---|
| Individual (N=993) | 0.594 |
| Season (N=6) | 0.015 |
| Cross-validation fold (N=10) | 0.016 |

| | |
|---|---|
| Residuals | 0.069 |

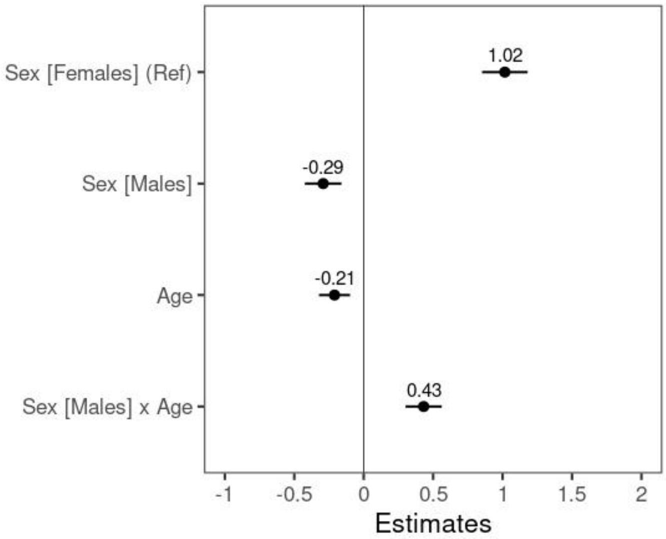


**Figure S1:** Coefficient plot of the estimates obtained from the mixed model (Table S2.B) testing the relationship between the scores predicted by the deep learning model and the age and sex of the individuals. Values correspond to the estimated mean and error bars to the 95% confidence interval in the estimated means.

#### Appendix 4: Visualisation of the features used for classification

**Table S3:** Between-sexes comparison of the proportion of activated pixels and within-regions mean activation of activated pixels. Comparisons were performed with multiple Student tests using 100 pictures of males and 100 pictures of females. For some pictures, some regions could not be analysed (due to the quality of the picture or errors in the automatic segmentation) and they were therefore removed from the analysis. “N females” and “N males” correspond to the number of individuals per regions used for the analysis. P-values were adjusted with the Benjamini-Hochberg procedure (Benjamini & Hochberg, 1995)

| | Region | N females | N males | Statistic | Adjusted p-value |
|---|---|---|---|---|---|
| Proportion of activated pixels | Beak | 100 | 100 | -25.70 | <0.001 |
| | Bib | 100 | 99 | -15.40 | <0.001 |
| | Cap | 100 | 99 | -14.00 | <0.001 |
| | Eyes | 94 | 93 | -14.40 | <0.001 |
| Mean activation of | Beak | 100 | 100 | -16.40 | <0.001 |

| | | | | | |
|---|---|---|---|---|---|
| the activated pixels | Bib | 100 | 99 | -3.54 | <0.001 |
| | Cap | 99 | 99 | -3.58 | <0.001 |
| | Eyes | 94 | 93 | -3.86 | <0.001 |

**Table S4:** Between-regions comparison of proportion of activated pixels and within-sex mean activation of activated pixels. Comparisons were performed with multiple Student tests using 100 pictures of males and 100 pictures of females. For some pictures, some regions could not be analysed and they were therefore removed from the analysis. P-values were adjusted with the Benjamini-Hochberg procedure (Benjamini & Hochberg, 1995).

| | Sex | Region 1 | Region 2 | Statistic | Adjusted p-value |
|---|---|---|---|---|---|
| Proportion of activated pixels | Female | Beak | Bib | -7.40 | <0.001 |
| | Female | Beak | Cap | -2.04 | 0.052 |
| | Female | Beak | Eyes | -2.27 | 0.018 |
| | Female | Bib | Cap | 3.98 | <0.001 |
| | Female | Bib | Eyes | 2.95 | 0.018 |
| | Female | Cap | Eyes | -1.08 | 0.285 |
| | Male | Beak | Bib | 1.40 | 0.166 |
| | Male | Beak | Cap | 3.70 | 0.001 |
| | Male | Beak | Eyes | 2.70 | 0.017 |
| | Male | Bib | Cap | 4.43 | <0.001 |
| | Male | Bib | Eyes | 1.77 | 0.095 |
| | Male | Cap | Eyes | -1.83 | 0.095 |
| Mean activation of activated pixels | Female | Beak | Bib | -5.21 | <0.001 |
| | Female | Beak | Cap | -0.68 | 0.498 |
| | Female | Beak | Eyes | 0.85 | 0.479 |
| | Female | Bib | Cap | 3.91 | <0.001 |
| | Female | Bib | Eyes | 4.65 | <0.001 |
| | Female | Cap | Eyes | 1.56 | 0.183 |
| | Male | Beak | Bib | 7.81 | <0.001 |
| | Male | Beak | Cap | 13.70 | <0.001 |
| | Male | Beak | Eyes | 10.60 | <0.001 |
| | Male | Bib | Cap | 5.23 | <0.001 |
| | Male | Bib | Eyes | 3.93 | <0.001 |
| | Male | Cap | Eyes | -0.71 | 0.480 |

#### Appendix 5: Distribution of age of the individuals in relation to sex

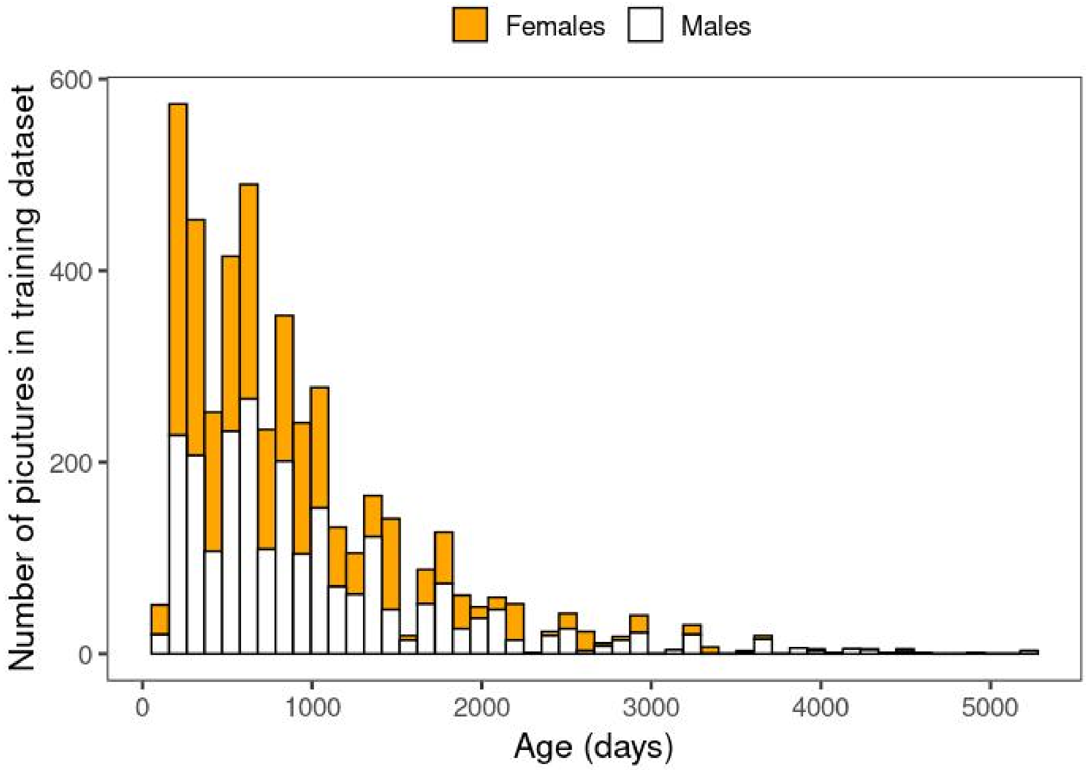


**Figure S2:** Stacked distribution of age of the individuals in function of sex in the training dataset. Females are in orange and males in white. Individuals over 3,000 days old are male-biased (29 females and 67 males).

**Table S5:** Estimate of the GLMM parameters to test the effect of sex and age on the score predicted by the deep learning network from pictures of the individuals' head, when excluding individuals older than 3,000 days. The model was computed 4,497 pictures.

| Fixed effect | Estimate | SE | 95% confidence interval | z-value | p-value |
|---|---|---|---|---|---|
| **Intercept** | 1.00 | 0.07 | 0.86 ; 1.15 | 13.62 | <0.001 |
| **Age** | -0.11 | 0.04 | -0.18 ; -0.03 | -2.76 | 0.006 |
| **Sex** - *Male* | -0.22 | 0.06 | -0.34 ; -0.11 | -3.76 | <0.001 |
| **Age** : **Sex** - *Male* | 0.36 | 0.05 | 0.26 ; 0.46 | 6.80 | <0.001 |

| Random effect | Variance |
|---|---|
| Individual (N=1,307) | 0.657 |
| Season (N=6) | 0.013 |
| Cross-validation fold (N=10) | 0.013 |
| Residuals | 0.060 |
